# The Role Of Senescent Cells Removal For Cell Culture Rejuvenation

**DOI:** 10.1101/2021.02.14.431016

**Authors:** Leonid Yu. Prokhorov

## Abstract

It is well known that aging reduces the number of healthy, vital cells, increases the number of dead cells, some of which are replaced by connective tissue. Similar processes occur in cell culture. If endlessly dividing transformed, or cancer cells grow in glass flasks and after their death for various reasons detach from the growth surface, lyzed and thus free up space for the growth of “young” cells, the culture lives arbitrarily long. But if in culture the “senescent” cells do not detach from the growth surface and do not make room for “young” cells, the “young” cells do not have the ability to devide and die. Because of this, the entire population of cells dies. Also in the body, “senescent” cells and intercellular tissues occupy a certain place in the body and do not allow “young”, “healthy” cells to divide.

## Introduction

Now it is clear that in fact there are no “eternal” cells, regardless of whether they are divided or not. Even in cultures of transformed or cancer cells, in which cells can divide an unlimited number of times, the latter are constantly dying. However, the place of dead cells occupy the “young”, daughter’s cells and therefore, this population lives for an unlimited time, subject to the constancy of habitat and the adequacy of nutrition. Therefore, it is necessary to speak not about “eternal” individual cells, and about of perpetual populations of cells [1].

Such “eternal” cell populations exist and are well known – these are cultures of transformed cells of different origins, different animals, as well as cultures of cancer cells of animals and humans. These cultures are constantly subcultivated in new flasks with a decrease in density, for example, 2 or 4 times, compared to the original flask. Then the cells in the new flask get a place to divide and the culture grows again to the so-called “saturating density” and the cells cease to divide as a result of contact inhibition.

However, even such “eternal” populations may die if they do not have space for reproduction and, accordingly, the ability to devide. This is a very important point and without a solution to this problem it will be impossible to achieve the goal of significantly increasing human lifespan [1].

Some authors believe that cell aging is one of the main paradigms of aging research. It began with the demonstration by L. Hayflick of a limited number of divisions of normal, untransformed cells that do not exhibit the properties of malignant cells [2], and these processes are largely regulated by the telomerase system [3]. In this case, the aging cell becomes the main actor in the aging process [4].

Since this initial discovery, the phenotypes associated with cellular senescence have expanded beyond growth arrest to include alterations in cellular metabolism, secreted cytokines, epigenetic regulation and protein expression. It is assumed that cellular senescence plays a role in brain aging and, notably, may not be limited to glia but also neurons [5].

Liu H. Y. and others [6] showed what can happen slowing aging animals (mice OVX-SAMP 8 with accelerated aging ovarectomy) due to the restoration of the “senescent” stem cells with platelet rich plasma.

Prattichizzo F. and others [7] note that aging is accompanied by a progressive decline of endothelial function and a progressive drift toward a systemic pro-inflammatory status. It is suggested that cell senescence may have a role in both processes and selective removal of senescent endothelial cells may hinder such harmful processes.

Other authors found that because, the accumulation of senescent cells in tissues is increasingly recognized as a critical step leading to age-related organ dysfunction then senescent vascular cells are associated with compromised bloodbrain barrier integrity [8].

Pluquet O. and other researchers [9] write that senescence is a complex cell phenotype induced by several stresses such as telomere attrition, DNA damage, oxidative stress, and activation of some oncogenes. It is mainly characterized by a cell enlargement, a permanent cell-cycle arrest, and the production of a secretome enriched in proinflammatory cytokines and components of the extracellular matrix. Senescent cells accumulate with age in tissues and are suspected to play a role in age-associated diseases.

Age-associated cardiovascular diseases are at least partially ascribable to vascular cell senescence. Replicative senescence and stress-induced premature senescence are provoked respectively by endogenous (telomere erosion) and exogenous (H_2_O_2_, UV) stimuli resulting in cell cycle arrest in G1 and G2 phases. Cell cycle analysis of endothelial cells transiently over-expressing ASncmtRNA-2 revealed an accumulation of endothelial cells in the G2/M phase, but not in the G1 phase [10].

When aging endothelial cells undergo senescence and manifest significant changes in their properties, resulting in impairment of the vascular functionality and neo-angiogenic capability. This leads to the emergence of several age-related diseases of the vascular system and other organs [11].

In order to show the need to create a permanent population of the existence of “young” cells in the body, experiments were conducted on cell cultures described in this paper.

## Materials and methods

The experiments use spontaneously transformed Chinese hamster cells (TCHC) obtained from subcutaneous connective tissue, line B11dii – FAF28, clone 431. Experimental cultures were grown in borosilicate glass Carrel flasks with a diameter of 54 mm hermetically sealed with rubber stoppers at a temperature of 37^0^C. For each experimental curve, 3 flasks were used. In an experiment started on 2.11.2002, the growth medium MEM with 10% or 1% of the bovin serum is used. TCHC is not subcultivated, i.e. the cells are not removed from the surface of the flask by ADTA and Tripsin treatmen, as in the classical cultivation, the flasks are not changed, but once in 15-16 days change the growth medium to fresh. Throughout the experiment since 2002, which continues to the present time, consider the number of living cells in each flask using an inverted microscope. We counted the number of cells in the field of view of the microscope on 5 sections of the flask and then calculated the number of cells per 1 square centimeter.

In another experiment, the same cells was used, but they grow on another growth medium – DMEM with 5% bovine serum with antibiotics (100 u/ml penicillin and 100 μg/ml streptomycin). The cells grew in the same borosilicate glass flasks without subcultivation and changing flasks and with the same periodic replacement of the growth medium. The experiment lasted equal 3.5 years (since 15.06.2013) and the cells live all the time in flasks without the classic subcultivation.

## Resalts

In figure 1 shows the change in the number of living cells in the culture of TCHC when periodic replacing the growth medium with 10% of bovin serum, but without changing the flasks. It can be seen that cells in the culture remain alive for more than 18 years (more than 6,700 days).

**Figure 1.**
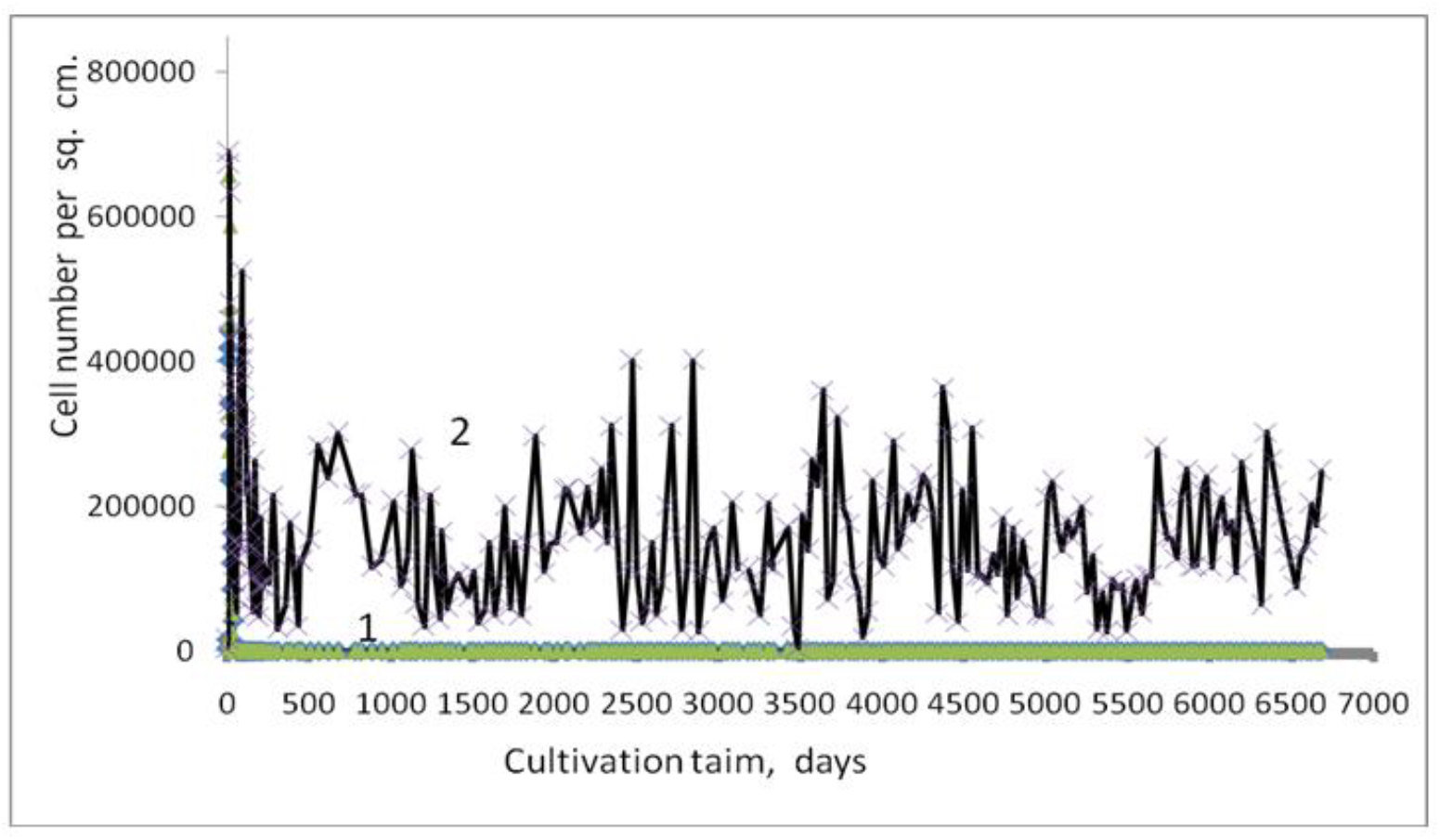
The experiment of 2002, the change in the number of living cells in culture growing on a medium with 10% bovine serum without changing the culture flasks: 1 – control – growth of the culture with one time changing the growth medium in 7^th^ day; 2 – the experiment – growth with periodic changing the growth medium.

In figure 2 shows the same experiment of 2002 initial period to 100 days the change in the number of living cells in culture without changing the culture flasks.

**Figure 2.**
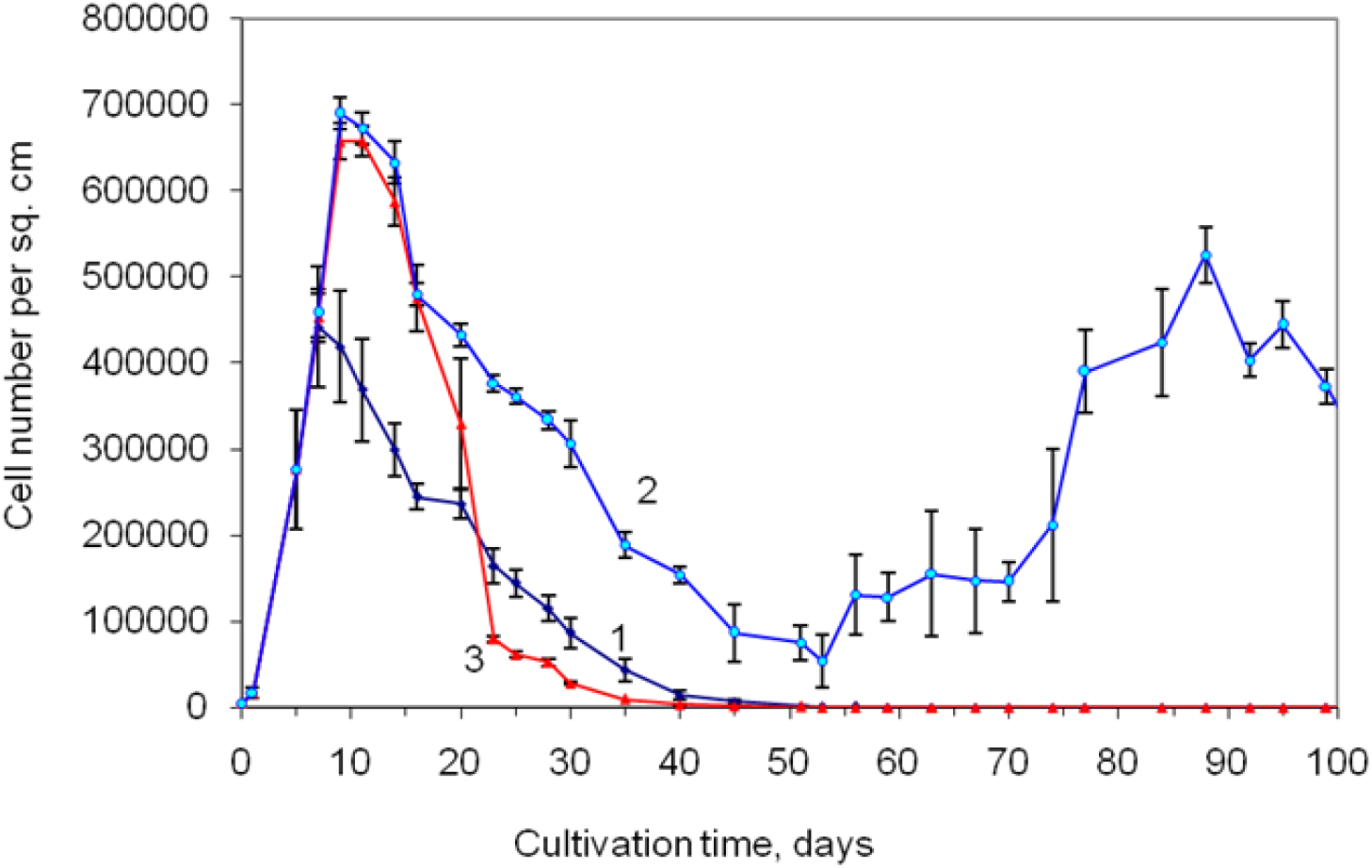
The same experiment of 2002 initial period to 100 days the change in the number of living cells in culture without changing the culture flasks:1- control – change in the number of living cells in the culture without changing the growth medium; 2 – the initial period of growth in the treated cell cultures with periodic replacement of the growth medium with fresh; 3 – the second control – change in the number of living cells in the culture with one changing the growth medium in 7^th^ day growth. On the chart indicate standard errors of the mean.

Figure 3 shows the type of cell culture in glass flasks at different times of growth without changing the growth medium: A – the first – second days after cell seeding at the beginning of the experiment or after replacing the medium with a fresh one later; B – 5-7 days of growth when the monolayer is reached; C – the aging period of a cell culture lasting 12-30 days after the cells have reached the monolayer and many cells die, and some still remain alive.

**Figure 3.**
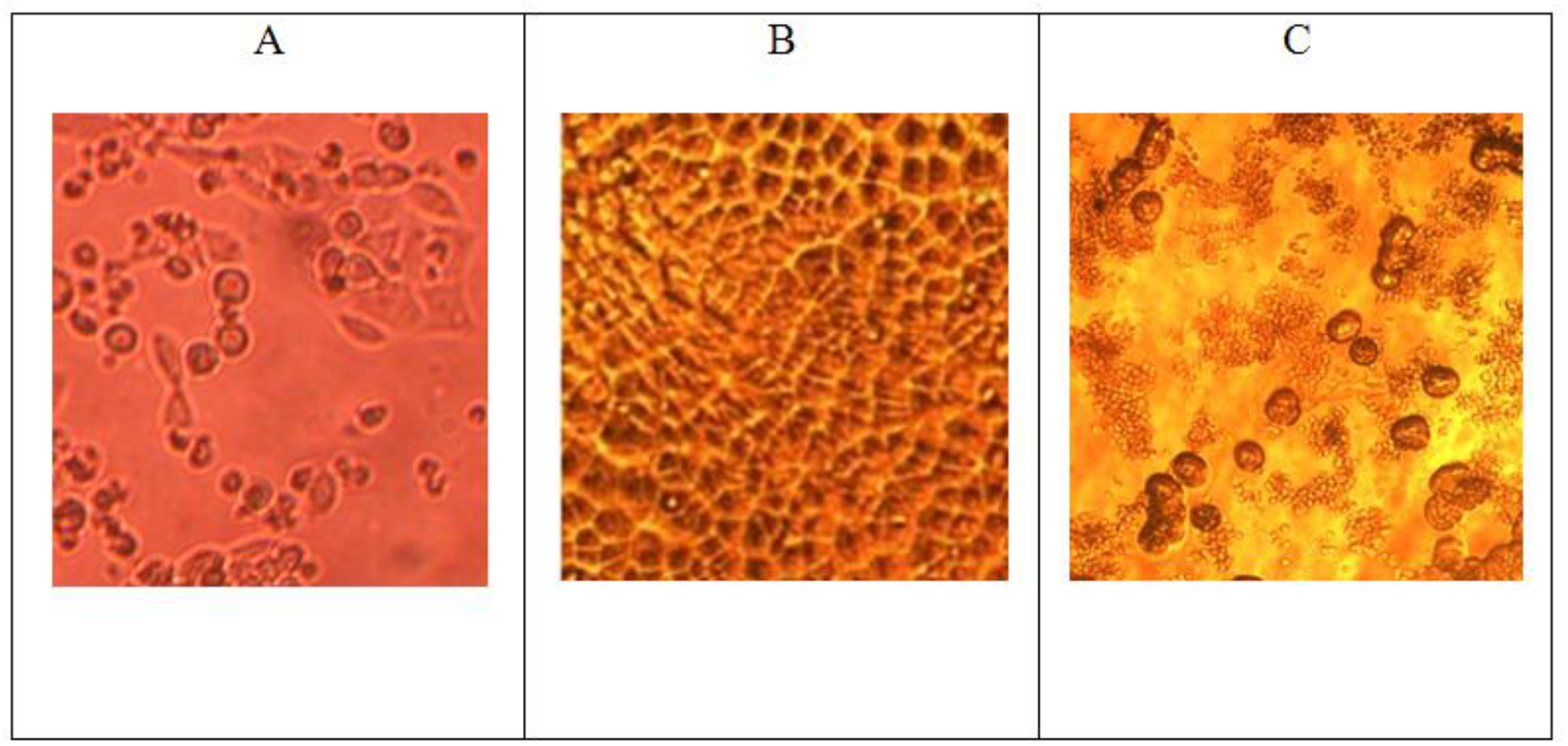
Type of cell culture in glass flasks at different times of growth: A – on the first day after the change meduam, B – after 5-7 days of growth when reaching the monolayer, C – during the aging culture on 12-30 days after reaching the monolayer, when many cells died, and some cells are still alive.

In control cultures (curve 1 on fig. 1, and curves 1 and 3 on fig. 2 and fig. 4) the growth meduam did not change or changed only once, so all the cells died after about 1 month.

**Figure 4.**
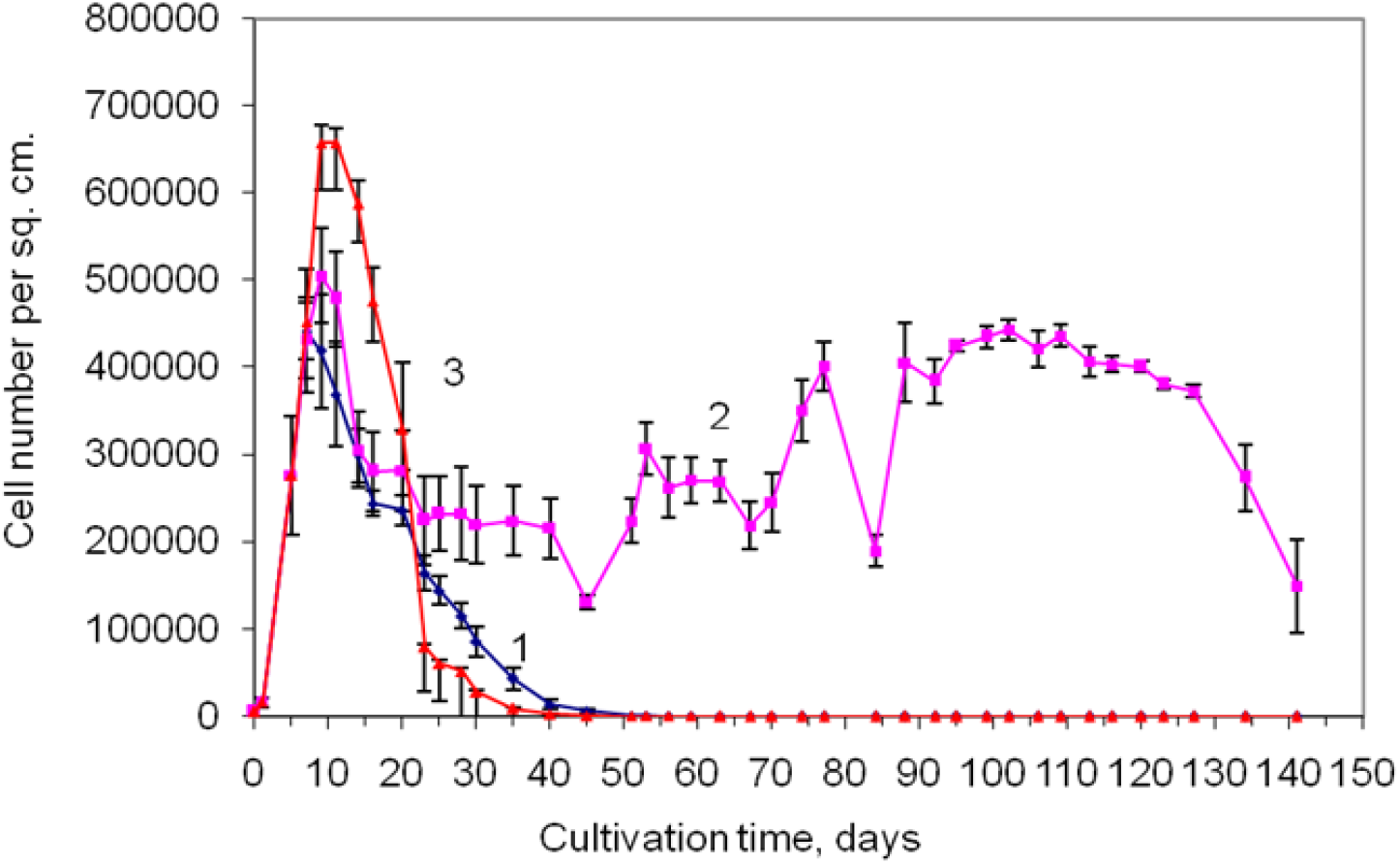
The change in the number of live cells in culture without changing the culture flasks: 1 – control – medium with 10% bovin serum did not change; 2 – periodical replacement of growth medium with 1% bovin serum starting from 7 days of growth after cell seeding; 3 – the second control – medium with 10% bovin serum replaced 1 time on the 7^th^ day of growth after cell seeding. On the chart indicate standard errors of the mean.

In the experiment (Fig. 1), when the culture reaches the monolayer, the “senescent” cells die, detach themselves from the growth surface and move to the growth medium. Thus they free up space for other cells. The remaining whole cells and the remnants of “senescent” cells are shown in figure 3C. If at this point to pour out the old growth medium, along with it “senescent” and destroyed cells are removed, and after adding a fresh medium, the remaining living cells in the flasks (“young” or “adult” cells – single cells in fig. 3C), begin are dividing again and again reach the saturating density (Fig. 3B). Then the process repeats itself.

In the 2013 experiment, when the growth medium with a bovine serum content of 5% is periodically replaced 2 times a month and without replacing culture flasks for 1267 days, the curve of change in the number of live transformed cells of the Chinese hamster is similar to the one shown in figures 1,2 and 4 for the 2002 experiment.

Thus, at concentrations of 1, 5 or 10% of bovine serum in growth meduam, the behavior of living cells in such a system with periodic replacement of the growth medium remains the same (Fig. 1,2 and 4).

It turns out that changes in the composition of the growht medium do not affect the general patterns of cell survival under the established conditions in accordance with the method. In these cultures, similar processes occur, that is, “senescent” cells die, breaking off from the growth surface, are transferred to the growth medium to make room for the growth of “young” cells. In the future, the “senescent” cells are removed from the flasks with the old medium, when it is replaced with a fresh one.

Unfortunately, the content of cultures with 1 and 5% bovine serum was discontinued for technical and financial reasons. However, it is obvious that the patterns of changes in the number of cells after the termination of experiments would be similar and further. Nevertheless, culture with periodic replacement of the growth medium with 1% bovine serum existed for 141 days and with 5% bovine serum – 1267 days, or almost 3.5 years, which is 4 and 42 times longer than the life span of the control culture.

These experiments also confirm the obvious need to remove “senescent” cells to make room for “young” cells so that the entire culture remains alive.

## Discussion

During aging, the number of healthy, vital cells decreases and the number of dead cells increases, some of which are replaced by connective tissue [12]. From this it follows: to solve the problem of aging, it is necessary that the cells of any organ always consist of viable, “young” cells capable to perform appropriate functions. The fact is that the “senescent” cells or intercellular tissue remaining instead of nonfunctioning “senescent” cells occupies a certain place in the body and does not allow the “young”, “healthy” cells to devide. In other words, “young” cells have no space for reproduction.

It also occurs in cell culture if the “senescent” cells do not detach from the growth surface and do not make room for “young” cells, the “young” cells do not have the ability to devide and die. Because of this, the entire population of cells dies [1, 13, 14, 15, 16].

Experiments are a convincing proof of the need to eliminate “senescent” nonfunctioning cells from the culture and, since cell cultures are a model that repeats patterns in the body [13], the results obtained on them can be transferred to whole organisms. In this case, the analogy lies in the fact, and in the culture and in the body over time, non dividing cells gradually die. Only in the body it is slow (for animals and people it is years and decades), and in culture it can be much faster (equivalent to a month). The study of long-term cultivation of cells in the described conditions allowed us to understand that for long-term preservation of cells in a viable state in the culture, it is necessary to eliminate the “senescent” cells, so that they free up space for “young” cells. Such spontaneous wave replacement of “senescent” cells with “young” cells allows the whole culture to live indefinitely. This same mechanism can be tried to reproduce in the whole body, if we want to ensure that the body is always dominated by “young” cells, not the “senescent” ones. It is the predominance of “young” cells over the “senescent” ones in any organ or tissue of the body that will ensure their viable functional state and, accordingly, the viability of the whole organism, overcoming aging and increasing life expectancy.

For cells to function properly and renew the population of organ or tissue cells, it is need to find a way to remove “senescent”, non-functioning cells from the organ or tissue.

If in culture the removal of “senescent” cells occurs by detaching them from the growth surface due to the weakening and destruction of protein bridges with the surface, in the body this process must be carried out forcibly. To remove “senescent” cells could try to use some enzymes that are able to destroy cell membranes. In particular, this could be the solutions of different concentrations of the enzymes trypsin, collagenase, elastase, chymotrypsin, hyaluronidase, bromelain, etc. [17].

Yosef R. et al. [18] studied the elimination of aging cells by inhibiting anti-apoptotic proteins BCL-W and BCL-XL. Joint inhibition of BCL-W and BCL-XL by siRNAs or the small-molecule ABT-737 specifically induces apoptosis in senescent cells. Notably, treatment of mice with ABT-737 efficiently eliminates senescent cells induced by DNA damage in the lungs as well as senescent cells formed in the epidermis by activation of p53 through transgenic p14(ARF). Elimination of senescent cells from the epidermis leads to an increase in hair-follicle stem cell proliferation. The finding that senescent cells can be eliminated pharmacologically paves the way to new strategies for the treatment of age-related pathologies.

Lutz H.U. [19] also tried to understand how “senescent” cells are eliminated from the body. He studied the elimination of “senescent” erytrocytes from the circulation as a result of the appearance of cell-age specific antigen on aging red blood cells. Selective phagocytosis of senescent human red blood cells requires a molecular alteration on the surface of aging red blood cells.

On the other hand Shimizu K. and Hokano M. [20] suggested that the disappearance of the cells containing iron in the red pulp was pointed out that “senescent” or worn out erythrocytes are eliminated more mechanical stress than the phagocytosis of macrophage of the spleen in patients with deposition of amyloid.

According to our assumption, to eliminate the “senescent” cells, you can try to apply ready-made preparations isolated from various plants or animals, for example, you can try to apply the already tested long-term protein extract from the snail Helix aspersa.

The feature of cell cultures survival due to the constant natural change of “senescent” cells by “young” ones described in this paper can be a significant addition to the general series of proofs for the theory of rejuvenation developed by the author and a significant increase in human lifespan [21].

This research did not receive any specific grant from funding agencies in the public, commercial, or not-for-profit sectors.

## Notes

### Competing Interest Statement

The authors have declared no competing interest.

## References

1. Prokhorov L Yu, Can aging be stopped? The reports of the Moscow Society of Nature Testers. Section of Gerontology. Given on 19 December 2003, Deposited in VINITI 11.10.2004 No. 1585. 2004, pp. 18–27 (In Rusian).

2. Hayflick L, The limited in vitro lifetime of human diploid cell strains. Exp. Cell Res. 1965; 37: 614–636. DOI: 10.1016/0014-4827(65)90211-9.

3. Olovnikov AM, A theory of marginotomy. The incomplete copying of template margin in enzymic synthesis of polynucleo tides and biological significance of the phenomenon. J. Theor. Biol. 1973; 41 (1): 181–190. DOI: 10.1016/0022-5193(73)90198-7.

4. Campisi J, Robert L. Cell senescence: role in aging and age-related diseases. Interdiscip. Top. Gerontol. 2014; 39: 45–61. DOI: 10.1159/000358899.

5. Tan FC, Hutchison ER, Eitan E, Mattson MP. Are there roles for brain cell senescence in aging and neurodegenerative disorders? Biogerontology 2014; 15(6): 643–660. DOI: 10.1007/s10522-014-9532-1.

6. Liu HY, Huang CF, Lin TC, Tsai CY, Tina Chen SY, Liu A, Chen WH, Wei HJ, Wang MF, Williams DF, Deng WP. Delayed animal aging through the recovery of stem cell senescence by platelet rich plasma. Biomaterials 2014; 35(37): 9767–9776. DOI: 10.1016/j.biomaterials.2014.08.034.

7. Prattichizzo F, Bonafè M, Ceka A, Giuliani A, Rippo MR, Re M, Antonicelli R, Procopio AD, Olivieri F. Endothelial cell senescence and inflammaging: microRNAs as biomarkers and innovative therapeutic tools. Curr. Drug. Targets 2016; 17(4): 388–397. DOI: 10.2174/1389450116666150804105659.

8. Yamazaki Y, Baker DJ, Tachibana M, Liu CC, van Deursen JM, Brott TG, Bu G, Kanekiyo T. Vascular cell senescence contributes to blood-brain barrier breakdown. Stroke 2016; 47(4): 1068–1077. DOI: 10.1161/STROKEAHA.115.010835.

9. Pluquet O, Pourtier A, Abbadie C. The unfolded protein response and cellular senescence. A review in the theme: cellular mechanisms of endoplasmic reticulum stress signaling in health and disease. Am. J. Physiol. Cell. Physiol. 2015; 308(6): 415–425.

10. Bianchessi V, Badi I, Bertolotti M, Nigro P, D’Alessandra Y, Capogrossi MC, Zanobini M, Pompilio G, Raucci A, Lauri A. The mitochondrial lncRNA ASncmtRNA-2 is induced in aging and replicative senescence in endothelial cells. J. Mol. Cell. Cardiol. 2015; 81: 62–70. DOI: 10.1016/j.yjmcc.2015.01.012

11. Regina C, Panatta E, Candi E, Melino G, Amelio I, Balistreri CR, Annicchiarico-Petruzzelli M, Di Daniele N, Ruvolo G. Vascular ageing and endothelial cell senescence: molecular mechanisms of physiology and diseases. Mech. Ageing Dev. 2016; 159: 14–21. DOI: 10.1016/j.mad.2016.05.003.

12. Strehler BL. Time, cells, and aging. New York and London: Academic press, 1962.

13. Prokhorov L Yu. Modeling of aging on stationary cell cultures. Dis... Cand. Biol. Sciences. Moscow, 1999 (In Rusian).

14. Prokhorov L Yu. Possibility increase life span of animals and man more than 1.5-2 times. Abstracts of the 10th Congress of International Association of Biomedical Gerontology. 19-23 September 2003, Cambridge, United Kingdom. Biogerontology 2003; 4 (Suppl 1): 1–107.

15. Prokhorov L Yu. Comparison of methods of replacement of “senescent” cells with “young” cells in the body and in the culture of cells growing without classical subcultivation. Clinical gerontology 2015; 21(11-12): 100–101 (In Rusian).

16. Prokhorov L Yu. The role of cell replacement techniques to increase the lifespan. Clinical gerontology 2016; 22(9-10): 62–63 (In Rusian).

17. Prokhorov LYu, Zgursky AA. Ways of the use of somatic and stem cells for the rejuvenation and treatment of human tissues and organs. The reports of the Moscow Society of Nature Testers (The 200th anniversary of the Founding of the Society). Moscow: LLC “Graficom-print”, 2005; 36: 119–121 (In Rusian).

18. Yosef R, Pilpel N, Tokarsky-Amiel R, Biran A, Ovadya Y, Cohen S, Vadai E, Dassa L, Shahar E, Condiotti R, Ben-Porath I, Krizhanovsky V. Directed elimination of senescent cells by inhibition of BCL-W and BCL-XL. Nat.Commun. 2016; 7: 11190. DOI: 10.1038/ncomms11190.

19. Lutz HU. Elimination of old erythrocytes from the circulation: exposure of a cellage specific antigen on aging erythrocytes. Schweiz. Med. Wochenschr. 1981; 111(41): 1507–1517 (In German).

20. Shimizu K, Hokano M. Elimination of old or worn red blood cells in the senile murine spleen. Acta Histochem. 1988; 83(1): 65–70. DOI: 10.1016/S0065-1281(88)80073-4.

21. Prokhorov L Yu. Is it possible to overcome aging? Today and tomorrow of cell therapy. Moscow: MAKS Press, 2017 (In Rusian).

